# Drp1 Proteins Released from Hydrolysis-driven Scaffold Disassembly Trigger Nucleotide-dependent Membrane Remodeling to Promote Scission

**DOI:** 10.1101/2025.04.10.648288

**Authors:** Elizabeth Wei-Chia Luo, Kelsey A Nolden, Haleh Alimohamadi, Michelle W. Lee, Liting Duan, R. Blake Hill, Gerard C. L. Wong

## Abstract

Dynamin-related protein (Drp1) drives mitochondrial fission, dysregulation of which leads to neurodegenerative, metabolic, and apoptotic disorders. The precise mechanism of fission completion is unclear. One prevailing model is based on GTP-driven *assembly* of Drp1 helices that increase confinement via force generation. However, constriction to nanoscopic tubule radii appears necessary but not sufficient for scission. The other is based on GTP-driven *disassembly* of a constricting Drp1 scaffold that drives a membrane disturbance, but the relation of disassembly to scission and GTP hydrolysis remain uncertain. Elucidation of mitochondrial fission is complicated by the multiple time-involved in the dynamics of mechanoenzyme activity, oligomer disassembly, and membrane remodeling. Using machine learning, synchrotron x-ray scattering, and a theoretical model, our data support a model where progressive GTP hydrolysis enable free Drp1s to increase their capacity for inducing membrane negative Gaussian curvature (NGC). Furthermore, we identify and Drp1 variants that diminish this progressive capacity. Machine learning reveals that predicted NGC-generating sequences of the Drp1 oligomer are not in contact with the confined lipid tube, that scission-enhancing membrane remodeling is triggered by free Drp1 released upon disassembly.

**Teaser:** Free Drp1 released from disassembly of oligomeric Drp1 constrictase promotes GTP-hydrolysis dependent mitochondrial scission.

## Introduction

The morphology of mitochondria and their intracellular size distribution are governed by a balance between mitochondrial fission and fusion (*1, 2*). Maintenance of this balance is crucial for normal cell function, and perturbations of this balance impact metabolism and apoptosis, and have been correlated to developmental defects, neurodegeneration, cardiovascular diseases, and cancer (*3*–*5*). Mitochondrial fission is the process by which a single mitochondrion divides into two daughter mitochondria, and is mediated by dynamin-related protein 1 (Drp1)(*6*–*8*), a GTPase from the dynamin superfamily. After three decades of work, it is generally agreed that Drp1 oligomerizes at the surface of mitochondrial membranes into helices with an outer diameter of ∼30nm (*9*), oriented with the GTPase domains facing the outside, and the variable domains (VD) on the inside, interacting with the mitochondrial membrane. Furthermore, there is also broad agreement that the Drp1 helix can undergo guanosine triphosphate(GTP)-dependent constriction to ultimately act as a GTP hydrolysis-driven effector of fission (*10*). Precisely how this happens remains a matter of debate, since constriction does not appear to lead deterministically to scission.

At present, there are two main classes of models for Drp1-driven fission and they start from radically different perspectives. In ‘two-step models’, Drp1 at a specific nucleotide loading state, initially forms a scaffold around the mitochondrion to create a confined lipid tubule. In the subsequent step, the GTP hydrolysis-triggered disassembly of Drp1 oligomers, which has been broadly observed experimentally (*11, 12*), leads to a disturbance which allows hemi-fission intermediates to form and thereby transition to a scission state. This model raises a number of questions. It requires Drp1 to have a high level of GTPase cooperativity, which is inconsistent with the measured Hill coefficient (*13, 14*). Moreover, a consideration of membrane viscoelastic relaxation times suggests that the lipid tubule should expand upon GTP-driven disassembly of the confining Drp1 oligomer, rather than scission. On the other hand, there are the ‘constrictase’ models, where Drp1 acts like a motor that enforces sliding of a helical turn, leading to increasing degrees of constriction. A number of questions are raised here also. It is known that *in vivo*, fission can be enforced by a single helical turn of Drp1 with 13-14 dimers. However, it is not known whether this arrangement will generate enough force or torque to constrict the tube, especially given that Drp1 itself needs to be progressively deformed. Also not entirely understood is the necessary dynamic coordination between different parts of the Drp1 structure for mechanical sliding, including the transient disruption of G-G domains and the states of hydrolysis at different positions. Finally, the relation of the motor activity to Drp1 disassembly is not known generally, although hypothetically, the hydrolysis-driven motor activity that leads to scission can eventually break the structure and lead to disassembly. These questions are complicated further by extreme changes in mitochondrial membranes during fission, from tubular structures with almost a constant mean curvature and zero Gaussian curvature (NGC) to catenoid-shaped necks in the irreversible hemi-fission state with zero mean curvature and strong NGC. The attending change in the metric imposes a high free energetic barrier in GTP-mediated constriction (*15, 16*). In the current state of knowledge, the relation of GTP-driven disassembly to scission is unclear. Ideally, seeing how Drp1 progressively mediates fission in a random, uncorrelated Drp1 population as a function of different ratios of GTP to guanosine diphosphate (GDP) hydrolysis *in vitro* can in principle be informative. However, most of the action in mitochondrial fission occurs well below the optical diffraction limit, and it is only quite recently that even the fundamental event of scission was demonstrated in narrow membrane tubules (*17*), a notable highlight after decades of work. The problem is further complicated by the different timescales involved in the Drp1-mediated fission process, from mechanoenzyme constriction activity, to hydrolysis-triggered oligomer disassembly, to membrane remodeling.

Here, we examine the action of a population of Drp1 on a model mitochondrial membrane at different effective ratios of GTP:GDP binding, using a combination of high-resolution SAXS, membrane mechanics calculations, machine learning, and mutation studies. Consistent with previous work on Dnm1, the yeast homolog of Drp1, we find that Drp1 is a GTPase that can also remodel membranes by inducing NGC, a saddle-shaped membrane deformation observed at scission necks during fission events (*18*). This indicates that there is underlying membrane remodeling capacity toward scission even for Drp1s. Using a validated machine learning classifier, we find that the NGC-generating regions in Drp1 are in the GTPase domain, which are distal to the Drp1 oligomer confined membrane tubule. This suggests that in the assembled Drp1 oligomer helix, these membrane remodeling regions are effectively turned off, and only turned on after hydrolysis-driven disassembly. Since we expect mixed Drp1 populations with different GTP hydrolysis states, it is important to assess not just how GTP hydrolysis influences membrane remodeling activity, but also how vulnerable this process is to heterogeneous hydrolysis states. These measurements are generally complicated by the dynamics of the evolving GTP:GDP ratio in an enzymatically active system. To obtain results with stable, well-defined GTP:GDP stoichiometries, in our experiments we replace GTP with a non-hydrolyzable GTP analog, GMP-PNP (henceforth denoted as ‘GTP(nh)’). Importantly, results here show that the change to GTP(nh) to GDP enforces a structural change in Drp1 with strong membrane remodeling consequences: Drp1-GDP induces drastically more NGC than Drp1-GTP(nh) on model mitochondrial membranes. We demonstrate that this change corresponds to a significant decrease in neck size upon GTP hydrolysis by using a mean-field calculation to extract mitochondrial neck sizes based on experimental SAXS data. Notably, we show that even heterogeneous mixed population of Drp1s with different ratios of Drp1-GDP to Drp1-GTP(nh) on model membranes progressively decrease neck radii via curvature-induced membrane remodeling. These results suggest that GTP hydrolysis not only releases free Drp1 from the assembled oligomers, but also drives Drp1 structural changes that actively add NGC to confined mitochondrial membrane tubules to enhance scission, via curvature-induced forces that act against membrane elasticity. In a companion paper, we show that the induced NGC by Drp1s can indeed trigger a ‘snap-through’ instability, constricting a lipid tube into narrow necks within the spontaneous hemi-fission limit (*19*). Interestingly, deletion of the membrane-interacting variable domain in Drp1ΔVD does not completely ablate its ability to induce NGC. The same is true for Drp1 G363D, a lethal assembly-deficient variant of Drp1. However, both variants lose the ability to cooperatively induce smaller necks with increasing Drp1-GDP/Drp1-GTP(nh) ratios. Taken together, these results highlight the role of free Drp1 released from GTP hydrolysis-triggered disassembly, and provide a point of contact between perspectives that have been difficult to reconcile in dynamin-like mechanoenzymes: GTP hydrolysis triggered disassembly releases Drp1s that induce NGC in sufficiently long lipid nanotubes generated by mechanoenzyme confinement, and precipitate a drastic “snapthrough” transition to produce scission.

## Results

### Machine learning predicts Drp1 to contain distinct membrane-remodeling elements

NGC, also known as saddle-splay curvature, exhibits concave curvature in one direction and convex curvature in the perpendicular direction. This specific type of curvature can be observed in membrane budding, membrane permeation, and cell division. Indeed, NGC is topologically required in neck regions during membrane fission and fusion events (*20*–*22*). Drp1 is a molecular motor that is known to promote membrane constriction to drive mitochondrial fission. Assessing the capacity of Drp1 to induce NGC, in addition to its mechanoenzyme activity, can shed light on how Drp1 influences mitochondrial remodeling.

To determine if Drp1 contains membrane-destabilizing subsequences that induce NGC in membranes, we utilized a support vector machine (SVM) machine-learning classifier designed to predict NGC-generating ɑ-helical peptide sequences (*23, 24*). This classifier was trained on membrane-permeating ɑ-helical antimicrobial peptides (AMPs), which have been experimentally observed to generate NGC in lipid bilayers by using SAXS (*25*–*27*). It effectively predicts whether a protein or peptide sequence has a high likelihood of inducing NGC. To screen potential NGC-generating subsequences in Drp1 (UniProtKB: O00429), a moving window scan of 10-35 amino acid length was performed. The machine-learning output scores, σ values, were calculated from inputting the window scan subsequences into the classifier. A σ score closer to +1 correlates with increased ability to induce NGC in membranes, whereas a σ score closer to 0 indicates low probability of membrane-disruptive activity. In order to identify Drp1 regions with high membrane activity, we aligned the window-scanned subsequences and took the normalized σ scores along the length of Drp1. (Fig. 1A) We identified three hotspots within the GTPase domain that have high σ scores, which indicated these subsequences have high probabilities for generating NGC on lipid membranes. To see the spatial relation between these NGC-inducing sequences, we colored the normalized σ scores on the crystal structure of Drp1 (PDB: 4BEJ). (Fig. 1B) Interestingly, the three NGC-generating ɑ-helices are clustered together on the side of Drp1 distal to the variable domain (VD), which is normally in contact with the confined mitochondrial membrane tube in the oligomeric form of Drp1. This structural arrangement implies that in the oligomeric form of Drp1, no NGC generation can take place because the helices are facing the wrong way. However, the oligomeric form of Drp1 in principle ‘stores’ NGC-generating capacity that can be liberated upon disassembly of the oligomer, which allows free Drp1 to rotate diffusely. In other words, Drp1 freed during oligomer disassembly, either in fragmented oligomeric form (e.g. dimer or tetramer) or monomeric form, can contribute to mitochondrial fission by inducing NGC in the confined membrane tube. These free Drp1s induced membrane deformations in mitochondrial model membranes are quantitatively measured in SAXS. Strong cubic phase signals are detected which indicated the Drp1-WT restructured membrane into phases rich in NGC, consistent with machine learning prediction (*19*).

**Fig. 1.**
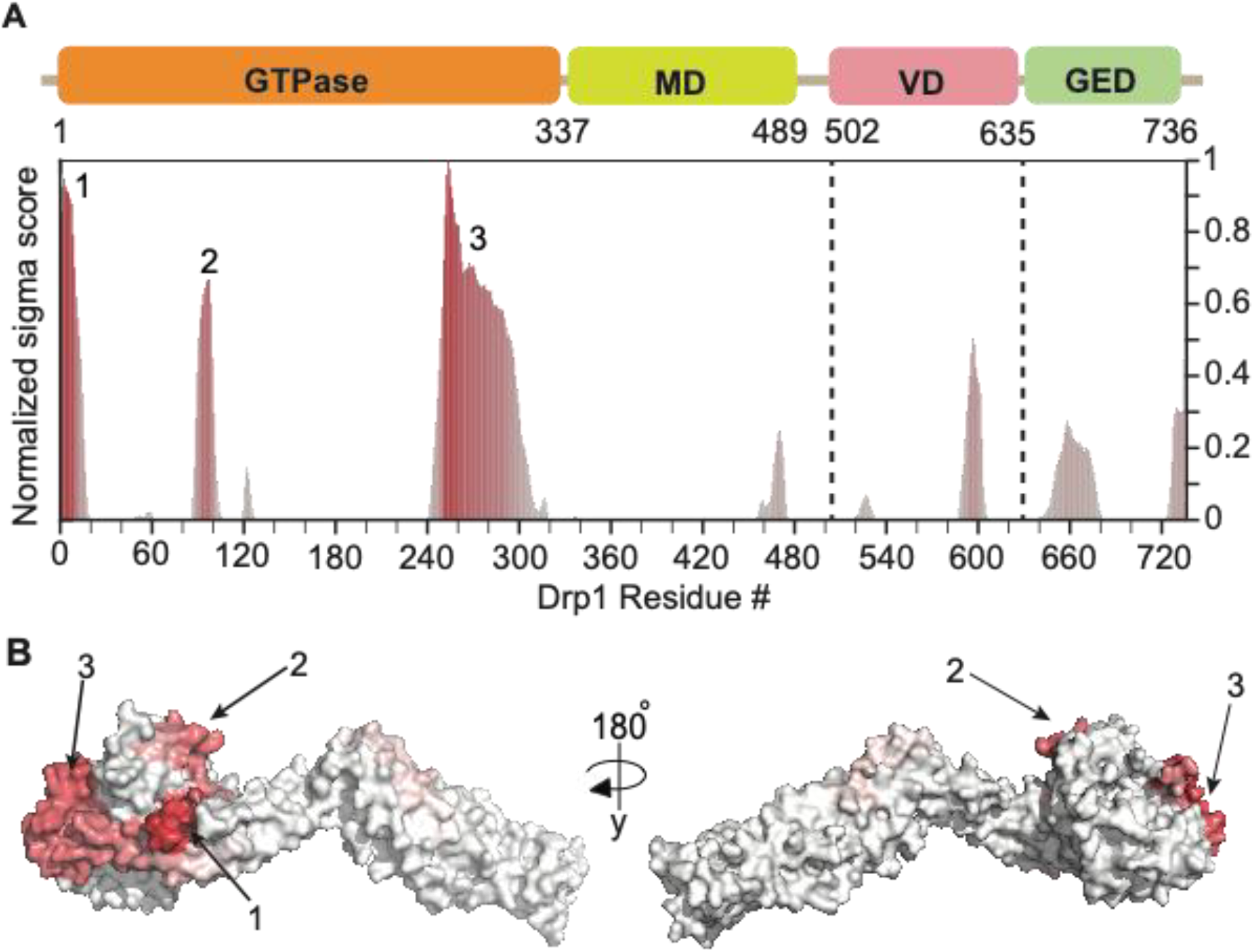
Drp1 has an intrinsic ability to induce negative Gaussian curvature and estimating the radius of mitochondrial scission neck based on the induced curvatures by Drp1 in the SAXS experiments. **(A)** A machine-learning classifier identifies regions within the Drp1 with high normalized σ scores of a moving-window scan. The top-scoring subsequences of Drp1 that may induce NGC are labeled with numbers. (SD: stalk domain, VD: variable domain, GED: GTPase effector domain) **(B)** 3D structure of Drp1 colored with normalized machine-learning σ score. (PDB: 4BEJ)

### Induced anisotropic curvature by GTP-driven disassembly of Drp1 helices can regulate mitochondrial fission

To estimate the size of the induced constricted neck by Drp1, we apply continuum elasticity theory to the SAXS measurements. We model the lipid bilayer as an elastic shell that can bend and adjust its configuration in response to imposed NGC. Assuming that the thickness of the membrane is negligible compared to bending (*28*), and in the limit of rigid proteins, the membrane-proteins interaction energy can be simplified using the generalized form of the Helfrich Hamiltonian energy given as (*28*–*33*)

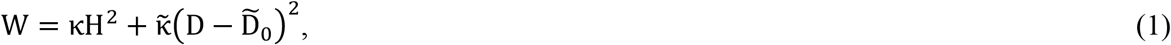

where κ and 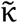 are the membrane bending rigidities, H is the mean curvature, and D is the curvature deviator (H^2^ − D^2^ =K is the Gaussian curvature) of the membrane. 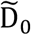 represents the induced deviatoric curvature by Drp1, which can be estimated from the cubic phases formed by Drp1 in SAXS given as 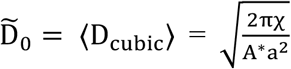 (*34*). Here, ⟨D_cubic_⟩denotes the average membrane curvature deviator in a cubic phase and a is the cubic lattice parameter. χ is the Euler characteristic and A^*^is the surface area per unit cell, which are specific to each cubic phase (*34, 35*).

Assuming the membrane is incompressible and in mechanical equilibrium, the normal force balance on the membrane surface gives the equilibrium shape of the membrane, and the tangential force balance results in the spatial variation of membrane tension (*33*). Having the amount of induced deviatoric curvature from SAXS measurements, we numerically solved the membrane shapes equations for a rotationally symmetric tubular membrane with a diameter of d^0^ =20 nm and a height to diameter ratio of L/d^0^ = 2 (*19*) (Fig. 2A). We assumed that 99% of the tubular membrane is covered by Drp1 (as shown in red in Fig. 2A) and set κ= 30K_B_T, where K_B_ is the Boltzmann constant and T is the temperature (*36*). In mechanical equilibrium and with no deviatoric curvature 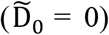, the membrane tension σ required to maintain a tubular membrane of diameter d_0_ is determined by minimizing the membrane bending energy with respect to d_0_, given as 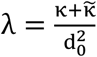(*37*). Membrane tension can vary in the order of O (0.1 pN/nm) (*15, 38*). In Fig. 2B, we plotted the radius of the constricted neck as a function of the Pn3m cubic lattice constant for different membrane tensions ranging from 0.09 pN/nm to 0.3 pN/nm.

**Fig. 2.**
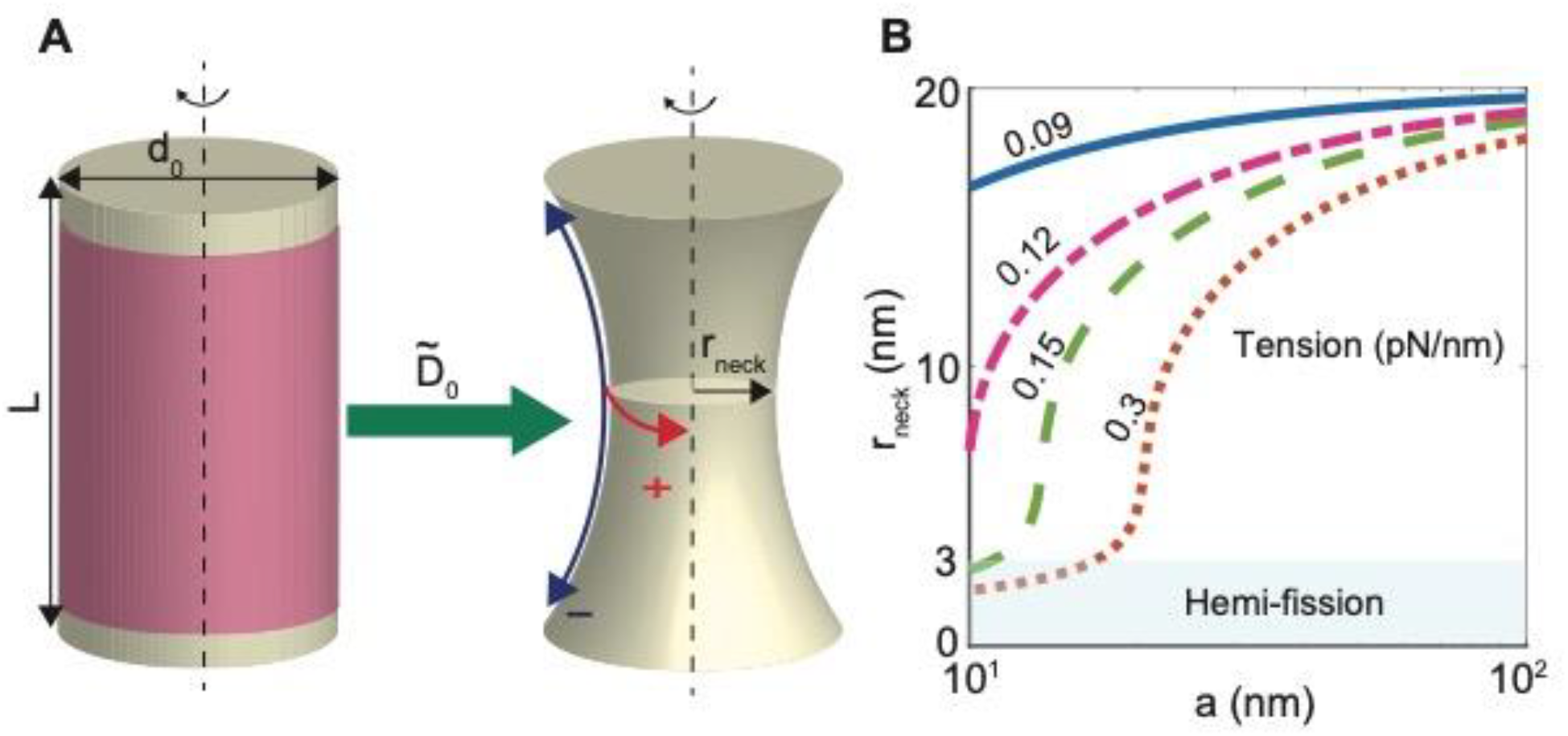
Estimating the radius of mitochondrial scission neck based on the induced curvatures by Drp1 in the SAXS experiments. **(A)** Remodeling of a tubular membrane with a diameter of d_0_ and a height of L/d_0_ = 2 into a constricted neck with a diameter of r_neck_ in response to the deviatoric curvature induced by Drp1. The yellow domain represents the bare membrane, and the red domain indicates the 99% area covered by Drp1. The structure and lattice constants of the *Pn3m* cubic phases formed in SAXS are used to estimate the magnitude of the induced deviatoric curvature 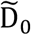 by free Drp1 proteins. The constricted neck is characterized by an NGC with opposite principal curvatures. **(B)** The radius of the constricted neck as a function of the *Pn3m* cubic lattice constant for different membrane tensions. The hemi-fission domain with r_neck_<3 nm is shown in blue.

Our results show that the deviatoric curvature induced by Drp1 can constrict a tubular membrane into a narrow neck (Fig. 2B). We also found that the degree of the neck constriction induced by the deviatoric curvature depends on the membrane tension (Fig. 2B). Higher membrane tension (λ >0.15 pN/nm) leads to a narrower neck (Fig. 2B). For example, at a small lattice constant of a= 10 nm, high membrane tension results in significant neck constriction, with a neck radius of r_neck_<3 nm, which is within the range of spontaneous hemi-fission (*39*–*41*). However, lower membrane tensions result in wider necks, with r_neck_ >8 nm (Fig. 2B). Consistently, previous studies have shown that high membrane tension is required for successful fission given the stochastic nature of the fission reaction with a low energy barrier transitioning from super-constricted to hemi-fission states. In vivo studies have also demonstrated that reduced membrane tension can delay fission from a few seconds to a couple of minutes (*15, 41*–*43*). Particularly, Morlot et. al., have shown the typical dynamin-induced fission time (a few seconds) corresponds to a high membrane tension of λ ∼ O (0.1 pN/nm) (*15*). From these results, we can conclude that a tighter curvature or a smaller lattice constant, as observed in SAXS measurements, corresponds to a narrower constricted neck, and higher membrane tension can facilitate the mitochondrial fission process. In the rest of the paper, we will use our mechanical model with high membrane tension of 0. 3 pN/nm to correlate the induced lattice constant in cubic phases to the radius of the mitochondrial constricted neck.

### GTP hydrolysis facilitates Drp1 mitochondrial remodeling activity

Similar to other large GTPases within the dynamin-related protein family, Drp1 is considered a mechanoenzyme that hydrolyzes GTP into GDP to enforce mitochondrial membrane scission (*17*). Machine learning results (Fig 1A) and SAXS experiments both consistently show that Drp1 has an intrinsic ability to induce NGC. It is interesting to assess how different nucleotide binding states affect the Drp1-mediated NGC generation since enzymatic activity is also an intrinsic dynamin function and can in principle influence its membrane remodeling capacity.

To investigate how the degree of GTP hydrolysis influences Drp1’s ability to constrict the fission neck, we incubated model mitochondrial membranes with Drp1 and either the non-hydrolyzable GTP analog Guanosine 5′-[β,γ-imido]triphosphate (GMP-PNP, denoted GTP(nh)) or GDP at various peptide/lipid molar ratios (P/L ratios). The mitochondrial membranes were modeled using ternary lipid mixtures of phosphatidylethanolamine (PE), phosphatidylcholine (PC), and cardiolipin (CL) at molar ratios of 75/5/20 and prepared as small unilamellar vesicles (SUVs).

In the SAXS spectra of SUVs with Drp1-WT, we observed a coexistence of phases: (1) an inverse hexagonal phase (H_II_) (peaks with q-ratios √1:√3:√4:√7:√9), (2) a bicontinuous cubic phase (Q_II_), and (3) a SUV form factor. (Fig. S1) In fact, there are two NGC-rich cubic phases commonly observed: The *Im3m* phase (peaks with q-ratios √2:√4:√6:√8) and the *Pn3m* phase (peaks with q-ratios √2:√3:√4:√6). The mitochondrial model membrane control sample of PE/PC/CL 75/5/20 SUVs only exhibited form factors. (Fig. S2A) The lattice constants for the cubic phases can be calculated from the linear regression fitting of the reflection peaks in the SAXS spectra. At P/L ratio 1/4000, Drp1-WT incubated with 1mM GMP-PNP showed an *Im3m* Q_II_ phase with the lattice constant of 42.309 nm. When the SUVs were co-administered with Drp1-WT and GDP, a *Im3m* Q_II_ phase is observed with the lattice constant of 35.9 nm. (Fig. 3A) Drp1-WT P/L ratio at 1/2000 was also tested and the same trend was observed. The *Im3m* lattice constants were 40.8 nm and 31.4 nm for samples incubated with GMP-PNP and GDP, respectively. (Fig. 3B) The SAXS spectra exhibited a mixture of H_II_ and Q_II_ phases, with cubic phase lattice constants decreasing from GMP-PNP to GDP at both P/L ratios. The smaller cubic phase lattice constant indicates a tighter membrane curvature with larger magnitudes of NGC. Using the cubic phase lattice constants as inputs, we estimated the radius of induced mitochondrial fission neck sizes using a continuum elasticity framework (*19*). Based on our results at the two P/L ratios 1/4000 and 1/2000, we find that the fission neck size changes from a larger diameter in the presence of GMP-PNP, to a smaller diameter in the presence of GDP. (Fig. 3C) This suggests that the Drp1 has a greater membrane curvature generating capacity after GTP(nh) has been replaced by GDP.

**Fig. 3.**
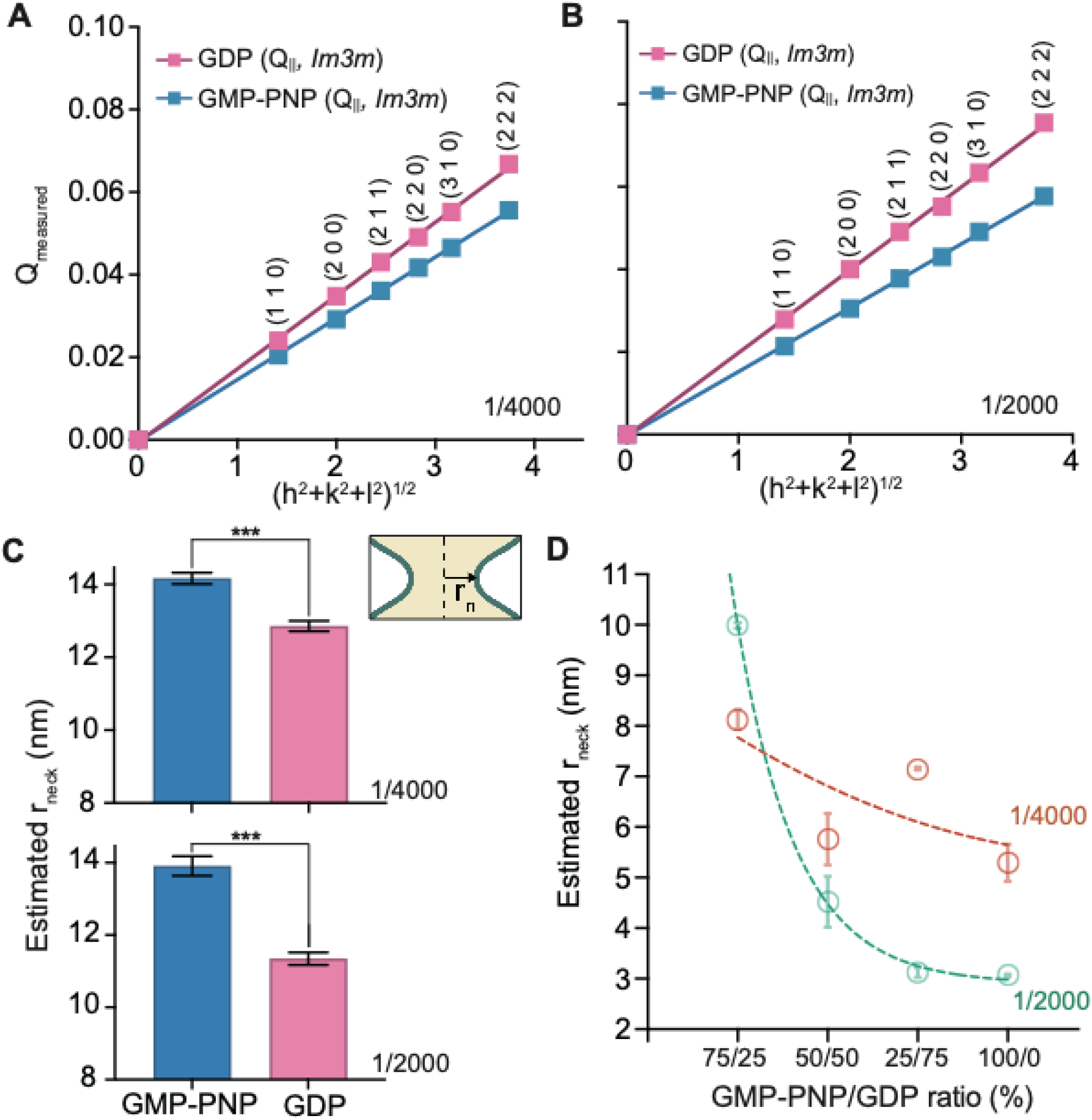
Nucleotide binding modulates Drp1’s ability to induce negative Gaussian curvature. **(A)** Indexing peaks from SAXS spectra of GDP or GMP-PNP introduced into the system with Drp1 and 75/5/20 PE/PC/CL model mitochondrial membranes. Plots of the measured Q positions, Q_measured_, versus the assigned reflections in terms of Miller indices. The lattice parameters were calculated from the slopes of the linear regressions. At P/L ratio 1/4000, sample at GMP-PNP binding state has a cubic phase lattice constant *Im3m* 42.5 nm. Samples treated with GDP have *Im3m* cubic phase lattice constants 35.9 nm. **(B)** At P/L ratio 1/2000, sample at GMP-PNP binding state has a cubic phase lattice constant *Im3m* 40.814 nm. Samples treated with GDP has *Im3m* cubic phase lattice constants 31.4 nm. **(C)** Plot the estimated fission neck radius (r_neck_) changes from the SAXS spectra in A. The estimated neck radius were calculated from the lattice constants. As Drp1 incubated with GMP-PNP or GDP, the neck radius (r_neck_) decrease. The same trend was shown in 2 P/L ratios: 1/4000 and 1/2000 **(D)** Titration in between 100% GMP-PNP and 100% GDP examine the GTP hydrolysis-dependent Drp1-WT neck size shrinkage. The estimated neck radius (r_neck_) were calculated from the lattice constants from the SAXS spectra.

Next, we investigated the neck sizes generated by heterogeneous Drp1-WT populations expected at progressively higher average levels of GTP hydrolysis. Notably, it is not a binary (‘first order’) transition between pure structural states of GTP(nh) and of GDP. A titration from GMP-PNP 100% to GDP 100% was performed at 25% increments. The equivalent mitochondria neck radii calculated from the SAXS data showed a monontonic composition-dependent decrease pure Drp1-GTP(nh) was mixed progressively more Drp1-GDP. At Drp1 P/L ratio of 1/2000, the calculated neck size shrank from 9.97 nm to 3.07 nm, 69.2% of decreasing in the radii. The same trend of gradual radii decrease (from 8.13 nm to 5.30 nm, 34.8% of decreasing) can also be shown at Drp1 P/L ratio 1/4000. (Fig. 3D) Moreover, at the higher P/L ratio 1/2000, larger neck size changes were observed, which indicated that higher Drp1 loadings drove larger neck size changes.

Taken together, our results suggest that Drp1’s intrinsic NGC induction can be modulated by the bound nucleotide. GTP hydrolysis enables Drp1 to generate a tighter curvature on the model mitochondrial membrane. A release of free Drp1s (shorter oligomers or monomers) can in principle impart membrane-remodeling activity to further constrict the mitochondrial fission neck, which is complementary to the GTPase hydrolysis-driven mitochondrial ‘constrictase’ activity. These findings suggest that free Drp1 released from disassembled oligomeric Drp1 helicoids can promote scission, and that even heterogeneous mixtures of Drp1 with different GTP hydrolysis states can enforce progressively narrower necks in a manner that depends on the average GDP content.

### Sub-functional Drp1 mutants retain ability to generate reduced NGC but completely lose the ability to monotonically decrease neck sizes with GDP/GTP ratios

So far, we have shown that Drp1 can precisely modulate fission neck size via membrane remodeling and do so in a manner that depends on GTP-hydrolysis state, with narrower fission necks as more Drp1-GTP(nh) is replaced by Drp1-GDP. Importantly, as the global population of Drp1 evolves from one rich in bound GTP(nh) to one rich in bound GDP, the system cooperatively and progressively enforces narrower fission neck sizes from the confined mitochondrial lipid tube. We examined how various mutations can dysregulate the mitochondria fission process in this context. The Drp1 variable domain (VD) has been proposed to be the site of interactions with lipids. However, the VD has been shown to be dispensable for both Drp1-mitochondrial recruitment and even some successful fission events, perhaps acting in an autoinhibitory manner (*44*). On the other hand, Drp1-G363D is a lethal point mutation of Drp1. This mutation allows the formation of tetramers but is impaired for higher-ordered assembly, which ultimately leads to the failure of mitochondrial neck fission (*45*). Here, we evaluate the ability of Drp1ΔVD and Drp1 G363D to cooperatively promote formation of fission necks as a function of increasing GTP hydrolysis (GDP/GTP ratio), and as a function of protein loading (P/L molar ratios).

Using SAXS, we examined SUVs made from model mitochondrial membranes incubated with Drp1ΔVD and Drp1 G363D at three P/L molar ratios (1/4000, 1/2000 and 1/1000). We find that the SAXS profile for Drp1ΔVD shows cubic phases at all three P/L ratios. With increasing P/L ratio, the cubic phase lattice constant decreases, which indicated there is dose-dependent curvature tightening in this process. The same trend was observed with Drp1 G363D. (Fig. 4A) Interestingly, these two mutations do not completely ablate the ability of Drp1 to induce NGC necessary for fission neck formation. To assess how mitochondrial fission may be disrupted by these Drp1 mutations, we tested how these two mutations (Drp1ΔVD and Drp1 G363D) behaved with heterogeneous Drp1 populations with different GDP/GTP ratios in terms of their ability to induce fission necks. (Fig. 4B). Like Drp1-WT, more protein led to more induced curvature. However, unlike Drp1-WT, Drp1ΔVD showed fission neck size increase between GMP-PNP and GDP at P/L 1/2000 (from 3.22 nm to 9.23 nm, 65.1% increasing) and 1/1000 (from 3.58 nm to 13.0 nm, 72.5% increasing). (Fig. 4B, left) For the Drp1 G363D mutation, we observe again that progressively larger Drp1 mutant protein loading on the membrane can lead to tighter curvatures, like WT. However, instead of inducing a tighter curvature when GTP(nh) is replaced by GDP, Drp1 G363D induced instead a subtle neck size *increase* or even no significant changes between GMP-PNP and GDP binding states. At P/L 1/4000 and 1/2000, the Drp1 G363D binding with GDP barely has no change on the size of the fission neck compared to G363D with GMP-PNP. (Fig. 4B, right) Only at highest P/L 1/1000, we observed an neck radii increasing from 3.93 nm to 7.79 nm (98.2% increasing). These results contrast markedly with WT behavior, where a significantly neck size decrease was observed when GTP(nh) was replaced by GDP in the protein. Both of these mutations disrupted the progressive monotonic decrease of mitochondria fission neck sizes for mixed Drp1 populations with progressively more bound GDPs.

**Fig. 4.**
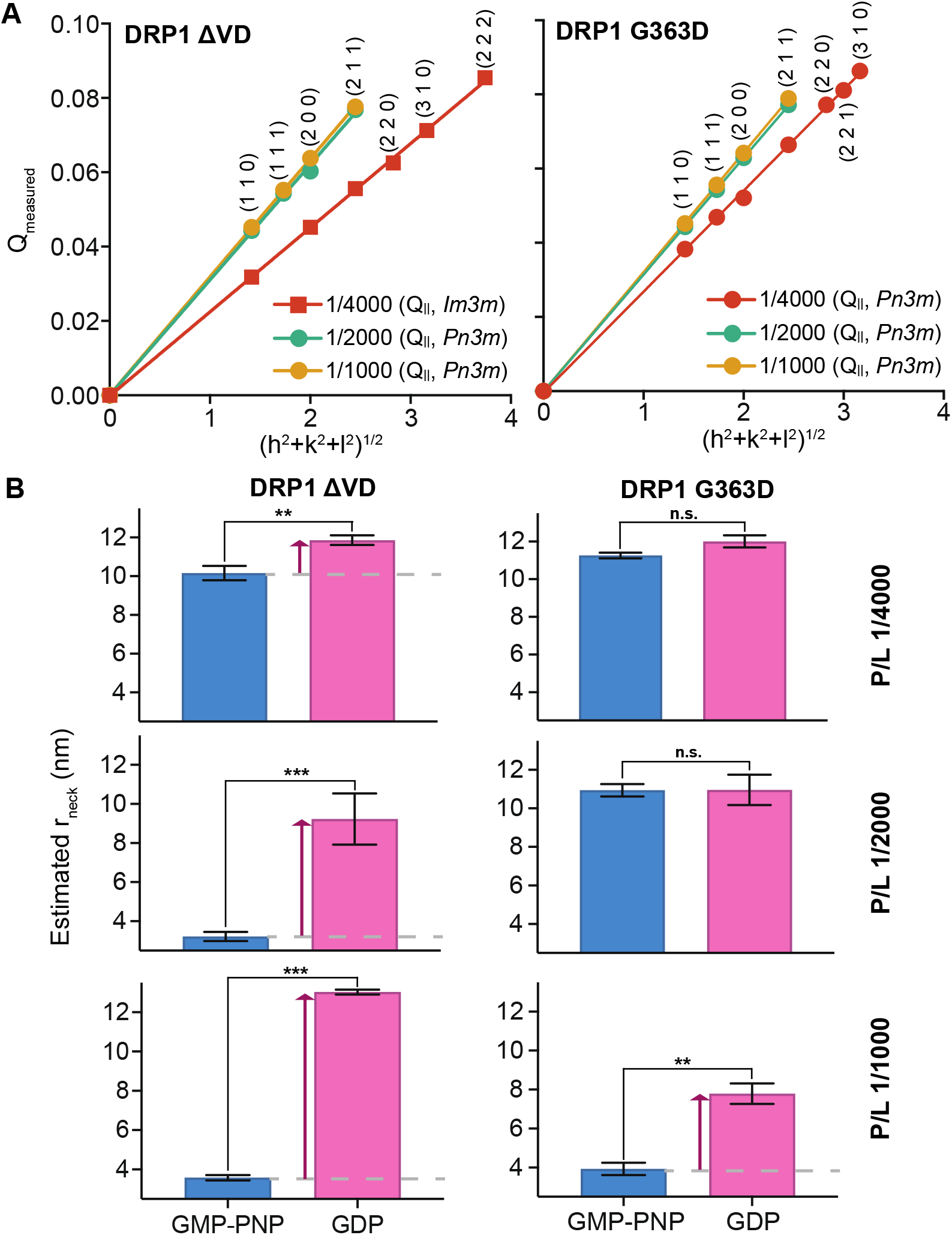
Sub-functional Drp1 mutants retain ability to generate reduced NGC but completely lose ability to control scission neck size with GTP/GDP ratios. **(A)** Drp1ΔVD (left) and Drp1 G363D (right) mutations restructured PE/PC/CL 75/5/20 model membranes into cubic phases. Indexing cubic phase peaks from SAXS spectra were plotted by measured Q positions, Q_measured_, versus the assigned reflections in terms of Miller indices. The lattice parameters were calculated from the slopes of the linear regressions. For Drp1ΔVD, the lattice constants for P/L ratio 1/4000, 1/2000 and 1/1000 are *Im3m* 27.8 nm, *Pn3m* 20.2 nm and *Pn3m* 19.7 nm respectively. For Drp1 G363D, the *Pn3m* lattice constants for P/L ratio 1/4000, 1/2000 and 1/1000 are 23.2 nm, 20.0 nm and 19.6 nm respectively. Both Drp1 mutations show the dose-dependent lattice constant decrease as P/L ratio increase. **(B)** Drp1ΔVD and Drp1 G363D mutations were incubated with the presence of GDP or GMP-PNP at P/L ratio 1/4000, 1/2000 and 1/1000. The estimated neck radii (r_neck_) calculated from the cubic phase lattice constants from panel A. Both Drp1 variants show the dysregulation of neck fission after the GTP hydrolysis. The arrows indicated the changes between the two nucleotide-binding states.

These findings above indicate that even the Drp1ΔVD and Drp1 G363D mutants remodel membranes by imparting NGC are consistent with our machine-learning based assessment of NGC-generating sequences in the mutant proteins. These Drp1 mutants still have intact NGC generating regions. More specifically, the NGC-generating hotspots and the mutation sites are located at opposite sides of the Drp1 3D protein. Because the hotspots remain intact in the mutants, their NGC-generating ability is expected to be preserved. These results suggest that the pathological mechanism of these Drp1 mutants correlate not with the loss of membrane remodeling activity, but rather with the loss of their ability to confine the lipid tube to the same degree, or their ability to maintain a strong monotonic decrease in the fission neck size as a function of average GDP/GTP ratio in the heterogeneous Drp1 population. These results also indicate that the activity of freed Drp1s, either in shorter oligomers or monomers, can be important for mitochondrial fission.

## Discussion

In this study, we determined that Drp1’s membrane remodeling activity unexpectedly differs depending on its nucleotide binding state. Our SAXS data showed that Drp1 itself possesses the ability to induce NGC on the membrane. Moreover, the transition from Drp1-GTP(nh) to Drp1-GDP leads to a progression from a low NGC state to a high NGC state, resulting in a decrease in the radius of the scission neck from large to small (see Fig. 2A). Thus, Drp1 with different bound nucleotides can cooperatively organize to generate fission necks with a size that depends monotonically on the GDP/GTP(nh) ratio, as evidenced by the decreased neck size observed in the GTP-GDP(nh) titration experiment (Fig. 3D). These new findings are consistent with a model in which Drp1 acts as a constrictase with the hydrolysis step generating disassembled Drp1 or significant reconfiguration of the Drp1 homooligomer that imposes NGC on the confined mitochondrial membrane tube, with GDP-bound Drp1 generating the most drastic changes. Curiously, deletion of the variable domain does not ablate the ability of Drp1 to induce NGC (Fig. 4A) despite its known role in binding cardiolipin-enriched membranes. Thus, Drp1 has intrinsic membrane activity independent of its variable domain. Our machine-learning classifier identifies the GTPase domain, which lie on opposite face of the Drp1 molecule as the variable domain, as a NGC-generating hotspot (Fig. 1A). This finding is particularly intriguing given that the GTPase domain is known to be in contact with the mitochondrial membrane during Drp1 helix pinching of the mitochondrion We also found that that the G363D pathogenic variant of Drp1 as well as the ΔVD constructs lose the ability to reduce fission neck size in a nucleotide-dependent manner (Fig. 4B).

In light of the ‘snap -through’ instability that can be induced by free Drp1 (described in our companion paper), the findings presented here lead to a new model for fission that reconciles previous contrasting perspectives based on constrictase activity and disassembly induced instability. Drp1 homooligomers generates narrow necks during the pre-constriction stage. In future steps, we propose that after the hydrolysis-driven disassembly of the confining Drp1 oligomer, then a free Drp1s (in shorter oligomeric form or monomers) population can undergo rotational diffusion to expose its NGC-inducing sequence motifs on the GTPase domain, thereby remodeling the membrane to drive the snap-through transition by inducing NGC. This remodeling event generates narrow necks in the pre-confined membrane tube, and can facilitate the hemi-fission state until complete separation occurs. Thus, scission is driven by the release of free Drp1 with GDP/GTP dependent membrane remodeling, a process that occurs in an environment with heterogeneous Drp1 populations with different GTP/GDP loadings. Moreover, since curvature inducing proteins are also curvature sensing proteins, our SAXS measurements suggest that free Drp1s exhibit a preference for adsorption onto highly curved membranes with a radius of curvature of around 10 nm, rather than onto the relatively large radii of mitochondria at 0.5-1 um (*46*). Therefore how effectively the released Drp1 precipitate the snap-through transition depends on the degree of membrane confinement from the initial constrictase activity.

In this study, we present a new framework for how Drp1 modulates the final step of mitochondrial fission. To build on our findings, it will be important to investigate this mechanism in a broader physiological context, particularly through in vitro and in vivo validation Particularly, mitochondrial fission is a highly coordinated process involving multiple proteins. For instance, during the pre-constriction stage, the endoplasmic reticulum–mitochondria encounter structure (ERMES) has been shown to regulate mitochondrial fission by coordinating the actions of actin, myosin, the endoplasmic reticulum, mitochondrial outer membrane receptors, and other associated proteins (*47, 48*). We envision that our findings can serve as a foundation for future studies exploring how these upstream players influence or integrate with the final membrane scission step mediated by Drp1.

Additionally, another intriguing direction for future research is the introduction of Drp1 mutations at the predicted NGC hotspots to experimentally validate our machine learning predictions in a cellular context. As a preliminary step, we performed an *in silico* analysis using our NGC machine learning classifier to identify potential mutation sites that could diminish NGC-generating ability (Sup. Fig. 4). By substituting each amino acid with all possible alternatives, we identified two key positions in two NGC-inducing helices as promising candidates (N9 and K10, and N254 and K255) (Sup. Fig. 4A, B). Interestingly, in the second helical structure of hotspot #3, no single point mutation ubiquitously reduced the predicted NGC score (Sup. Fig. 4C). These findings provide a foundation for designing Drp1 variants to further test our proposed mechanism.

We believe that the findings presented here provide a point of contact between the almost diametrically opposing premises in our models for mitochondrial fission, by showing how disassembly of the oligomer can contribute to generation of scission necks in a counterintuitive manner, specifically, by releasing free Drp1s with membrane remodeling activity to drive a snap-through transition in the confined membrane tube. Furthermore, we showed how Drp1 mutants can dysregulate this process and potentially lead to pathological clinical consequences.

## Materials and Methods

### Machine-based learning classifier and 3D modeling

The sequence of human Drp1 isoform 1 (UniProt: O00429) was screened against a machine-learning classifier to predict protein subsequences with NGC-inducing membrane activity using a window scanning size of 10-35 amino acids(*49*). For each computed subsequence, the classifier calculates a prediction score (σ) and a probability (P(+1)) of NGC inducing activity. The σ scores were normalized for each amino acid using the sum of all the σ scores for subsequences containing that amino acid, divided by the number of subsequences containing that specific residue. A heatmap of the NGC hotspot prediction scores (σ) was generated by plotting the normalized σ scores against the sequence of Drp1 whereby σ scores closer to 1 are considered as having a higher probability to generate NGC. The σ scores were then displayed in a white-to-red gradient, representative of low to high σ, on the x-ray crystallography structure of human Drp1 (PDB: 4BEJ). Regions of beta-sheet secondary structure predicted to have NGC-inducing activity were excluded moving forward as the machine learning classifier was trained on alpha-helical peptide sequences.

### Neck size estimation

For the given Helfrich energy in Eq. 1, in mechanical equilibrium, and a rotationally symmetric coordinate, the normal and tangential force balances on the membrane simplified to a system of first order differential equations given as

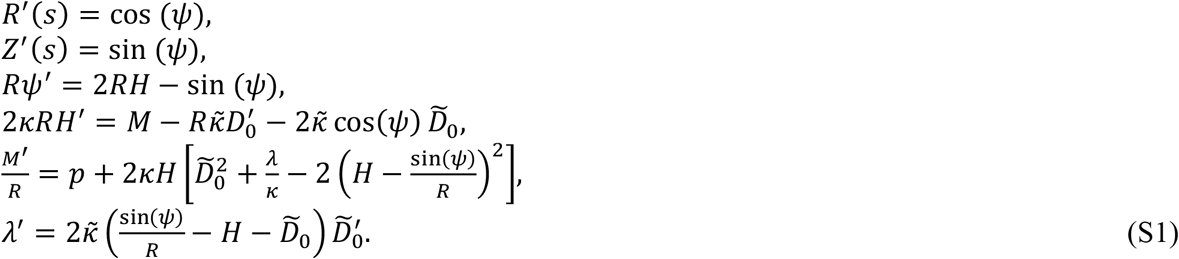

where *s* is the arc-length along the membrane, *R(s)* is the radial distance from the axis of rotation, *Z(s)* is the elevation from the reference plane, *ψ* is angle made by the tangent with respect to the horizontal, and (·)^′^ is the first order derivative with respect to the arc-length. To solve the coupled system of equations, we prescribe the following boundary conditions at the two ends of the simulation domain

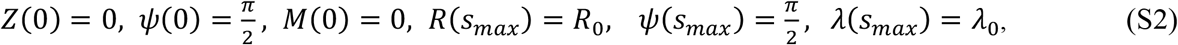

where *s*_*max*_ is the maximum length of the computational domain and *λ*_0_ is the prescribed membrane tension at the boundary. We used a boundary value problem solver in MATLAB (“bvp4c”) to solve the system of equations (S1) coupled with the prescribed boundary conditions (S2). le. We refer the interested reader to our companion paper (*19*) for details on the mathematical principles underlying this model.

### Drp1 size calculation

The size of Drp1 in nanometers was estimated using the previously solved and deposited x-ray crystallography and cryo-electron microscopy structures of Drp1 (PDB: 4BEJ and 5WP9, respectively). Each structure was imported into the molecular modeling program PyMOL (Schrödinger) and the linear distance between distal GTPase domains of two protein monomers within a dimeric unit, the proposed base functional unit of Drp1, was calculated using the Measurement Wizard tool in angstroms. Measurements were then converted to nanometers using the conversion factor of 1 Å = 0.1 nm. We estimated *R*_*protein*_ ∼ 7 nm based on the 2D projected area of the Drp1 structure on the surface.

### Drp1 purification

Recombinant human Drp1 isoform 1 was expressed in *Escherichia coli* as a TEV-His 6 fusion construct and purified using nickel-affinity chromatography as previously described (*50*). Harvested cells from 1-liter cultures were lysed using an EmlusiFlex C3 homogenizer (Avestin) at 15,000 p.s.i. The resulting cell lysate was clarified by centrifugation (15,000 rpm, 4 °C, 45 minutes) using a JA-20 fixed-angle rotor in a Beckman J2-21 centrifuge. The clarified cell lysate supernatant was applied to a 5 mL nickel affinity column (GE Healthcare; Sepharose high performance beads) equilibrated in Buffer A and purified by FPLC prior to elution with a high imidazole buffer. Fractions containing eluted protein were pooled and dialyzed into 20 mM Hepes, pH 7.4, 500 mM KCl, 0.1% BME overnight at 4 °C using 6-8 kDa molecular weight cut-off dialysis tubing (Repligen). The expression tag was concurrently removed during dialysis using recombinant TEV protease (∼1:20 ratio mg/mg). Remaining protease and affinity tag were then removed using a reverse nickel chromatography step. Purified protein was concentrated using a 30 kDa spin-concentrator and then dialyzed once more using 6-8 kDa molecular weight cut-off tubing into Drp1 buffer (20 mM Hepes, pH 7.4, 150 mM KCl, 2.0 mM MgCl2, 1 mM DTT). Protein concentration was determined using the theoretical extinction coefficient and protein molecular weight. Purified protein was flash-frozen in liquid nitrogen in single-use aliquots and stored at -80 °C until use.

### SAXS sample preparation and data collection

Lyophilized phospholipids 1,2- dioleoyl-sn-glycero-3-phosphocholine (DOPC), 1,2-dioleoyl-sn-glycero-3-phosphoethanolamine (DOPE), and cardiolipin (CA), purchased from Avanti Polar Lipids, were dissolved in chloroform at 20 mg/mL to produce individual lipid stock solutions. Ternary lipid compositions were prepared from these stock solutions as mixtures of DOPE/DOPC/CA in a molar ratio of 75%/5%/20%, respectively. The ternary mixes were thenevaporated under nitrogen and desiccated overnight under vacuum to form dry lipid films. Lipid films were resuspended in Drp1 buffer (20 mM N-(2-hydroxyethyl)piperazine-N’-ethanesulfonic acid (HEPES), pH 7.4, 150 mM KCl, 2 mm MgCl_2_, 1 mM Dithiothreitol (DTT)) to a final concentration of 20 mg/mL. Lipid suspensions were incubated overnight at 37 °C, sonicated until clear, and extruded through a 0.2 μm pore Anotop syringe filter (Whatman) to yield SUVs. SUVs were then mixed with recombinant protein(Drp1, Drp1 G363D, or Drp1ΔVD) at specific P/L molar ratios as indicated. For nucleotide-containing samples, Guanosine 5’-diphosphate (Sigma-Aldrich, GDP) and Guanosine 5’-[β,γ-imido] triphosphate (Sigma-Aldrich, GMP-PNP a non-hydrolyzable analog of GTP) were added at a final concentration at 1 mM. Precipitated protein– lipid complexes were transferred into 1.5 mm quartz capillaries (Hilgenberg GmbH, Mark-tubes) and hermetically sealed with an oxygen torch. SAXS measurements were collected at the Stanford Synchrotron Radiation Lightsource (SSRL) (beamline 4-2) using monochromatic X-rays with energies of 9 keV. Samples were incubated at 37 °C and centrifuged using a tabletop centrifuge before measurement. Scattered radiation was collected using a DECTRIS PILATUS3 × 1M (with a 172 μm pixel size). The resulting 2D powder diffraction patterns were azimuthally integrated into 1D patterns using the Nika1 1.76 package for Igor Pro 7.04 (Wavemetrics). For all samples, multiple measurements were taken at different times to ensure consistency.

### SAXS data analysis

SAXS data analysis was performed as described previously (*20, 25*). Briefly, to determine the lipid phases present in each sample, the integrated scattering intensity (I(q)) was plotted as a function of scattering intensity (q) using MATLAB (MathWorks). The q-values corresponding to peak positions were determined and their ratios were compared to the permitted reflections for different liquid-crystalline lipid phases (e.g., lamellar, hexagonal, cubic). Lamellar phases exhibit integer ratios of 1:2:3 and hexagonal phases exhibit ratios of √1:√3:√4:√7:√9:√12:√13. Cubic phases observed in our experiments belonged to the Pn3m space group, which permits reflections at ratios of √2:√3:√4:√6:√8:√9, and the Im3m space group, which permits reflections at ratios of √2:√4:√6:√8:√10:√12:√14:√16. For each cubic phase, the measured peak positions were related to the Miller indices (h, k, l) of their observed reflections with the equation q = 2π√(h^2^+k^2^+l^2^)/a, where a is the lattice parameter. The experimentally determined q-values were plotted against √(h^2^+k^2^+l^2^) and fit to a linear regression model. The resulting slope of the line was then used to calculate a. The average Gaussian curvature ⟨K⟩ per unit cell volume for a cubic phase was calculated using the equation ⟨K⟩=2πχ/A_0_a^2^, where χ is the Euler characteristic and A_0_ is the surface area per cubic unit cell. For Pn3m cubic phases, χ = −2 and A_0_ = 1.919. For Im3m cubic phases, χ = –4 and A_0_ = 2.345. To estimate the neck size, the lattice constants of the cubic phases in the SAXS spectra were converted into neck sizes based on our mean field model. To estimate the error and get the error bars, the range of cubic lattice constants was converted into a range of estimated neck radii. The significance between two data points was determined by calculating the highest confidence interval percentage at which their error ranges do not overlap.

## Author contributions

Conceptualization: GCLW, RBH, EWL, KAN, HA

Methodology: GCLW, RBH, EWL

Formal analysis: EWL, HA

Funding acquisition: RBH, GCLW

Investigation: EWL, HA

Project administration: GCLW, RBH

Resources: EWL, HA, KAN, GCLW, RBH

Software: EWL, HA Supervision: GCLW, RBH

Validation: EWL, GCLW

Visualization: EWL, HA

Writing—original draft: EWL, HA, KAN, GCLW, RBH

Writing—review & editing: EWL, HA, ML, LD, GCLW, RBH

Co-first authors are ranked in the order in which they participated in the experiment.

## Data and materials availability

All data needed to evaluate the conclusions in the paper are present in the paper and/or the Supplementary Materials.

