## Supplementary Materials for "Drp1 Proteins Released from Hydrolysis-driven Scaffold Disassembly Trigger Nucleotide-dependent Membrane Remodeling to Promote Scission"

Elizabeth Wei-Chia Luo *et al.*

**This PDF file includes original SAXS spectra:**

Figs. S1 to S4

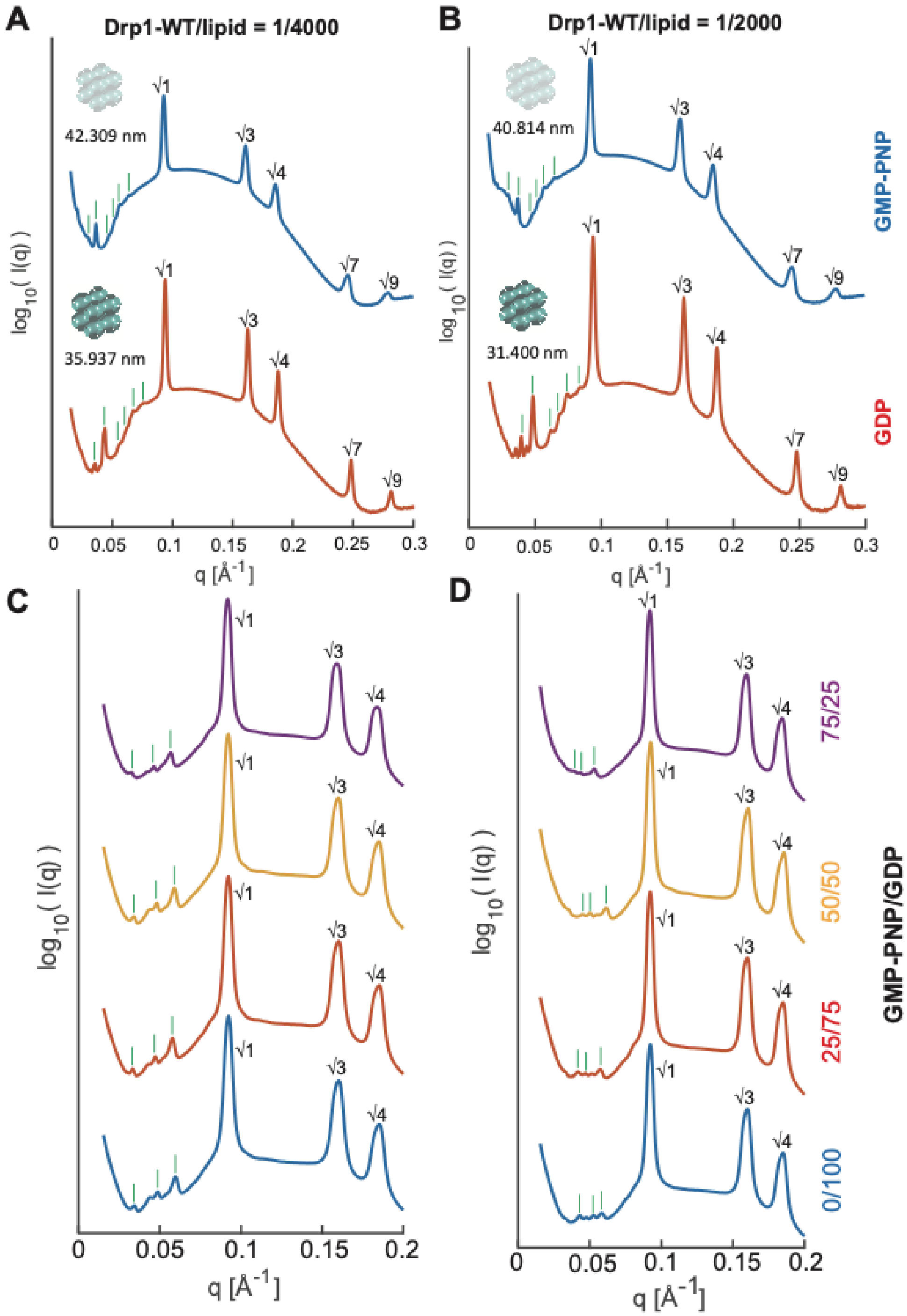


**Fig. S1.**

**Original SAXS spectra of model mitochondrial membranes incubated with Drp1-WT at different nucleotide-binding states. (A and B)** Drp1-WT incubated with PE/PC/CL 75/5/20 model membrane at 1/4000 (A) and 1/2000 (B) protein/lipid (P/L) ratios. The 3D representations of Im3m cubic phases were presented in the islet with the lattice constants aside. Samples are treated under 100% GMP-PNP (blue) or 100% GDP (red) conditions. **(C and D)** Drp1-WT mixed with lipid membranes under varies GMP-PNP/GDP ratios. P/L ratio 1/4000 (C) and P/L ratio 1/2000 (D) were both tested. The GMP-PNP/GDP ratios are: 75/25 (purple), 50/50 (yellow), 25/75 (red), and 0/100 (blue) to mimic the interstates from GTP hydrolyzed to GDP. **(A-D)** Observed reflections for the Pn3m or Im3m cubic (green) and hexagonal (black) phases have been assigned on the curves.

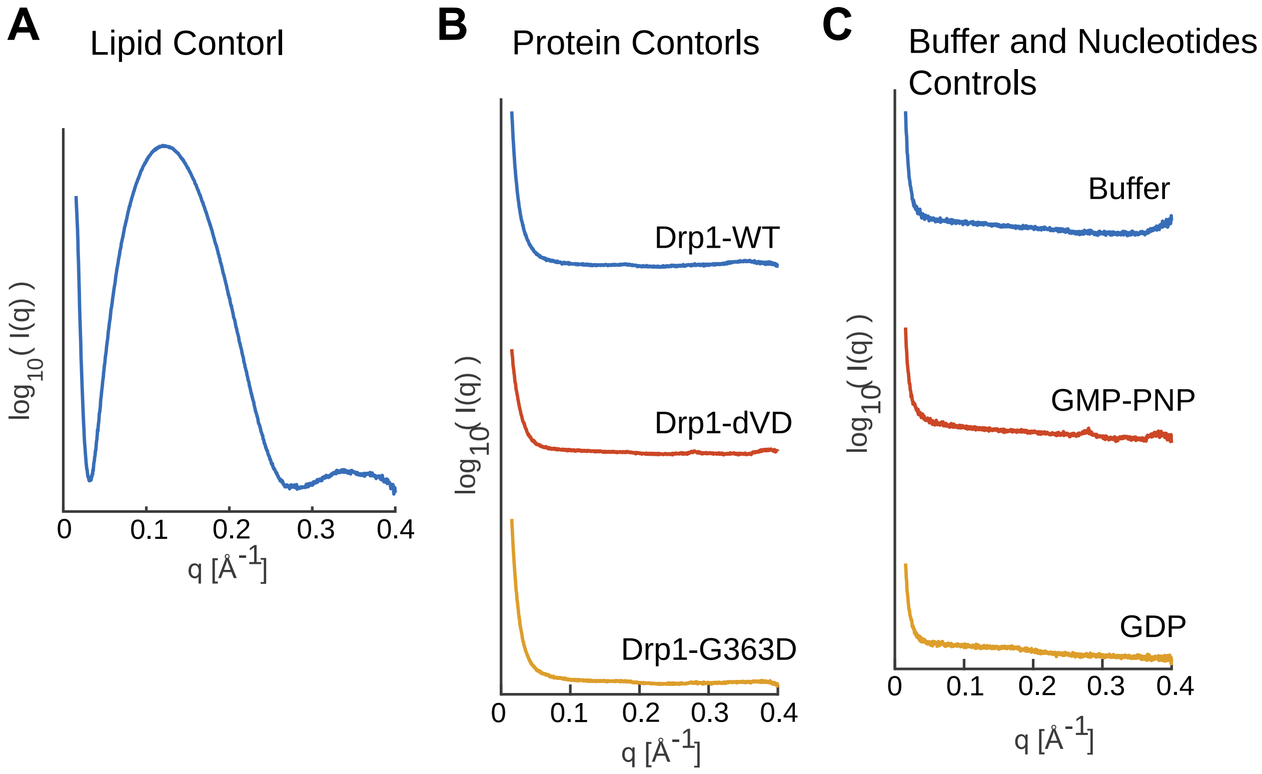


**Fig. S2.**
**Original SAXS spectra of the control experiments. (A)** The lipid (PE/PC/CA 75/5/20) only control. **(B)** Protein only controls for the 3 Drp1s tested in this work. **(C)** pH7.4 buffer and the nucleotide solution only controls. Only form factors identified in the spectra.

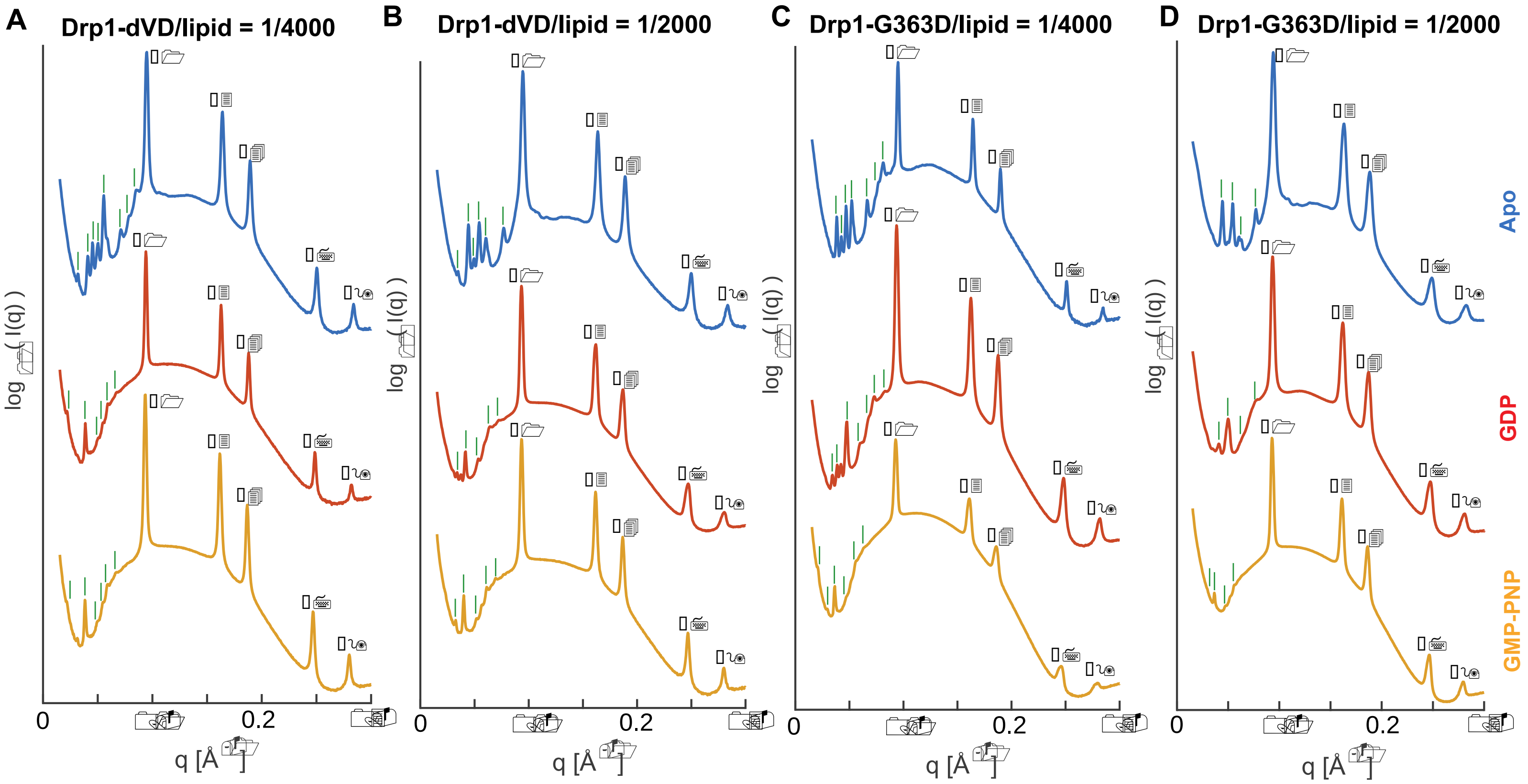


**Fig. S3.**
**Original SAXS spectra of model mitochondrial membranes incubated with Drp1-ΔVD and Drp1-G363D at different nucleotide-binding states. (A and B)** Drp1-ΔVD incubated with PE/PC/CL 75/5/20 model membrane at 1/4000 (A) and 1/2000 (B) P/L ratios. **(C and D)** Drp1-G363D mixed with lipid membranes at 1/4000 (C) and 1/2000 (B) P/L ratios. **(A-D)** Drp1 mutants and the lipids were incubated under no nucleotides (blue), 100% GDP (red) or 100% GMP-PNP (yellow) conditions. Observed reflections for the Pn3m or Im3m cubic (green) and hexagonal (black) phases have been assigned on the curves.

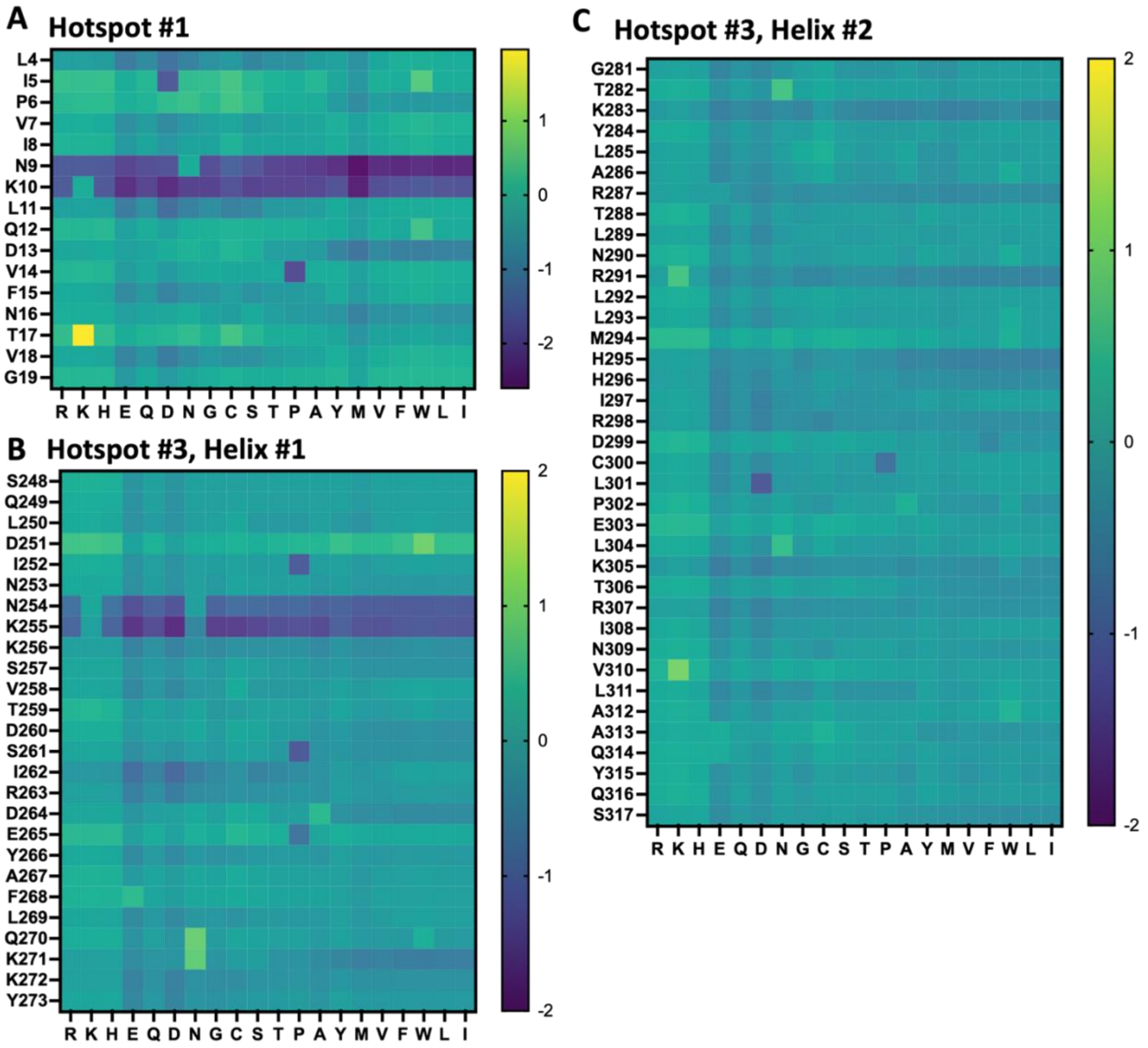


**Fig. S4.**

**2D heat map of the membrane activity fitness landscape of the Drp1 NGC hotspots. There are three helical regions within the NGC hotspots identified in the Figure 1A from the main text. (A-C)** We conducted machine-learning screens to each helical regions with various point mutations. Fitness of each unique single mutants is calculated as $\Delta\sigma = \sigma\text{mutant} - \sigma\text{WT}$. Mutations predicted to reduce membrane activity are shown in dark blue, while those that increase membrane activity are shown in yellow. **(A)** Hotspot #1, in main Fig. 1A, is an α-helical structure. N9 and K10 point mutations strongly reduce the NGC prediction score, making them good mutation candidates for reducing membrane remodeling activity. **(B and C)** There are two helical regions within hotspot #3, in main Fig. 1A, and the machine-learning screening were performed separately. **(B)** N254 and K255 point mutations in helix #1 strongly reduce the NGC prediction score, making them good mutation candidates. **(C)** In helix #2, none of the mutation sites consistently reduced the NGC prediction score across all substitutions. Only C300P and L301D showed some potential for lowering NGC scores.
